# A digital collection of rare and endangered lemurs and other primates from the Duke Lemur Center

**DOI:** 10.1101/688739

**Authors:** Gabriel S. Yapuncich, Addison D. Kemp, Darbi M. Griffith, Justin T. Gladman, Erin Ehmke, Doug M. Boyer

## Abstract

Scientific study of lemurs, a group of primates found only on Madagascar, is crucial for understanding primate evolution. Unfortunately, lemurs are among the most endangered animals in the world, so there is a strong impetus to maximize as much scientific data as possible from available physical specimens. MicroCT scanning efforts at Duke University have resulted in scans of more than 100 strepsirrhine cadavers representing 18 species from the Duke Lemur Center. Scans include specimen overviews and focused, high-resolution selections of complex anatomical regions (e.g., cranium, hands, feet). Scans have been uploaded to MorphoSource, an online digital repository for 3D data. As captive (but free ranging) individuals, these specimens have a wealth of associated information that is largely unavailable for wild populations, including detailed life history data. This digital collection maximizes the information obtained from rare and endangered animals without degradation of the original specimens.

## Introduction

Lemurs, a radiation of primates endemic to Madagascar, are an important group of animals for understanding the evolutionary history and adaptive origins of primates. Unfortunately, they are among the most endangered mammals in the world, with 94% of lemur species threatened by extinction [1]. Continued habitat degradation and fragmentation, illegal poaching, and challenging economic and political circumstances in Madagascar mean that lemurs are likely to remain under acute threat in the foreseeable future [1]. While conservation groups have developed several local, site-specific action plans [2], protecting and studying these animals requires multiple strategies both in Madagascar and internationally.

The Duke Lemur Center (formerly the Duke University Primate Center) is a prime example of an alternative approach to the conservation and scientific study of lemurs. Founded in 1966 in Durham, North Carolina, the Duke Lemur Center (DLC) was established to operate as a “living laboratory” and permit non-invasive study of rare primates, including galagos, lorises, and lemurs (which together comprise the primate suborder Strepsirrhini). Over its history, the DLC has housed more than 4,000 individuals from 39 primate species, and currently houses nearly 240 individuals from 17 strepsirrhine species. The DLC is involved in multiple lemur conservation efforts, including 1) managing several breeding populations as an Association of Zoos and Aquariums accredited institution; 2) the SAVA conservation program, a community-based approach to sustainable forest management and economic improvement in northern Madagascar; and 3) working with the Malagasy government to develop animal husbandry, welfare, and breeding programs for ex-situ lemur populations in Madagascar.

While the cofounders of the DLC, Dr. John Buettner-Janusch and Dr. Peter Klopfer, ran research programs focused on genetics and behaviour respectively, the DLC’s unique resources have provided data for a wide variety of scientific fields, including biomechanics, physiology, and palaeontology. The importance and rarity of the animals housed at the DLC necessitates thorough and effective use in educational and research initiatives, and this spirit of efficiency extends to treatment of deceased individuals. When an animal dies at the DLC (most frequently of old age), veterinary staff perform necropsies to remove internal organs and then preserve cadavers in cold storage for future research purposes. There are currently more than 400 cadavers in storage. However, because DLC cadaveric specimens are available for destructive sampling, the total information preserved by each specimen decreases over time. Digitizing the DLC’s cadaveric collection presents an opportunity to preserve hard tissue data without degrading specimen integrity, thereby maximizing the educational and scientific value of these rare and endangered animals.

Here we present an open access 3D digital collection of microCT scan data representing 113 adult individuals from the Duke Lemur Center. The data presented here were generated by GSY and ADK for use in their dissertations [3,4] and the collection will continue to grow with future scanning efforts. All scans are publicly available on MorphoSource.org [5], an online repository specifically designed to archive 3D data. Primate cadavers have long been recognized as valuable scientific resources that could be utilized much more efficiently [6,7], provided scientists motivated by different research questions could coordinate specimen access. The digital collection presented here follows several recent efforts to digitize and publish unique and valuable datasets [8–10].

## Materials and Methods

### DLC cadaveric collection

A total of 483 microCT (μCT) scans of strepsirrhine primates housed at the Duke Lemur Center were performed at the Shared Materials Instrumentation Facility at Duke University. Specimens represent both major clades of strepsirrhine primates, including 82 lemurs, 17 galagos, and 14 lorises (Figure 1). Among these individuals, two lemurs and five loris specimens were iodine-stained to permit visualization of soft tissue anatomy. Due to physical space limitations of the μCT machine, the sample is biased toward species less than 3 kg; larger species housed at the DLC (e.g., *Propithecus* and *Varecia*) have not yet been scanned, though methods developed in the course of this study will allow them to be scanned in the future. Additional demographic information for most specimens is available in Zehr et al. [11]. Table 1 provides a summary of the sample by species, and individual specimens, scanning parameters and DOIs for all currently available data are provided in Table 2.

**Fig 1.**
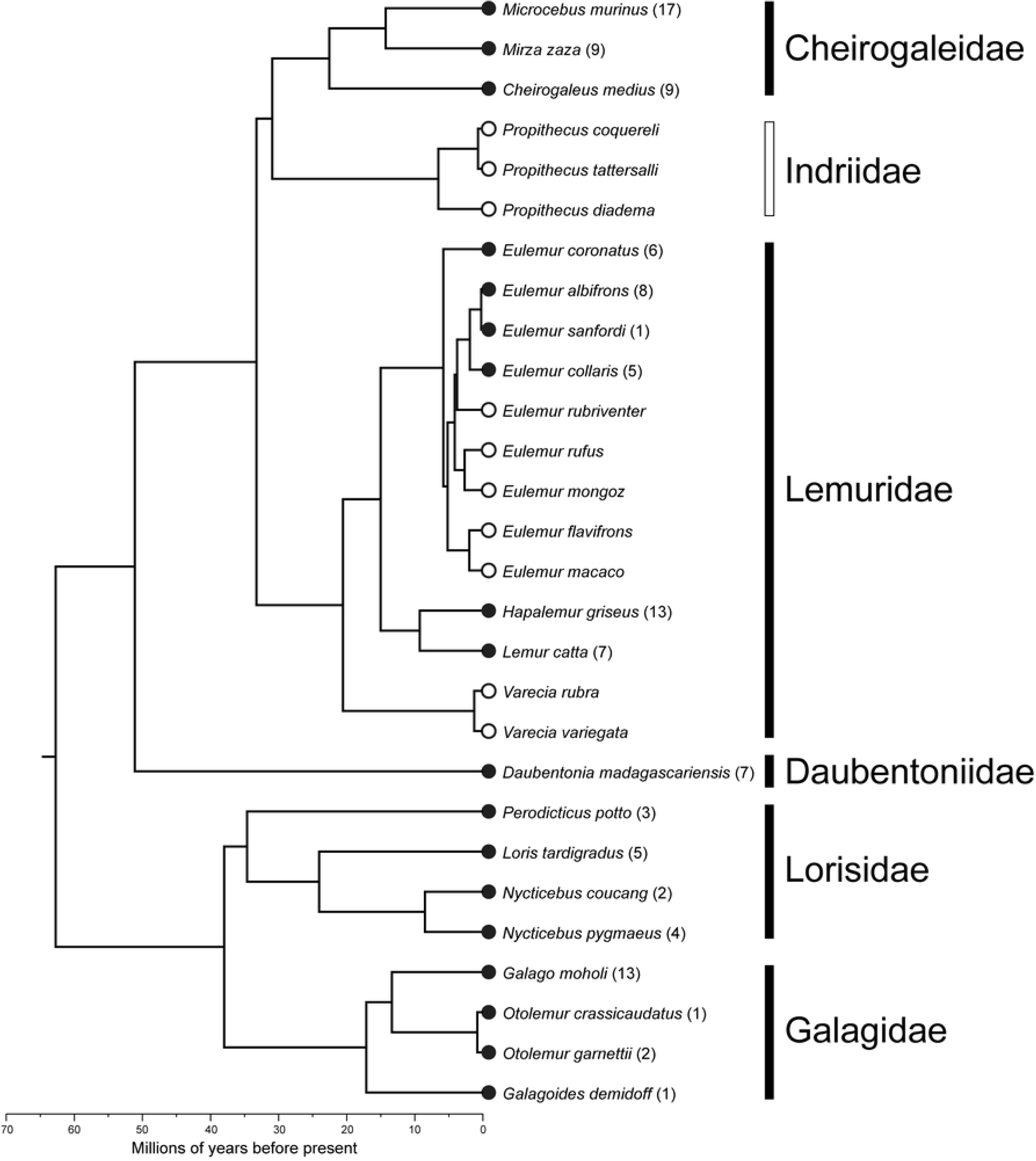
Strepsirrhine phylogeny and number of individuals included in this microCT collection. Dark circles and bars indicate taxa currently represented in the collection. Open circles and bars represent taxa housed at DLC but not currently scanned. Chronogram downloaded from 10ktrees.org, version 3 [16].

**Table 1.** Summary of sample by species.

| Species | *n* individuals | *k* scans |
| --- | --- | --- |
| *Cheirogaleus medius* | 9 | 35 |
| *Daubentonia madagascariensis* | 7 | 35 |
| *Eulemur coronatus* | 6 | 23 |
| *Eulemur albifrons* | 8 | 31 |
| *Eulemur collaris* | 5 | 18 |
| *Eulemur sanfordi* | 1 | 5 |
| *Galago moholi* | 13 | 53 |
| *Galagoides demidoff* | 1 | 4 |
| *Hapalemur griseus* | 13 | 57 |
| *Lemur catta* | 7 | 35 |
| *Loris tardigradus* | 5 | 23 |
| *Microcebus murinus* | 17 | 72 |
| *Mirza zaza* | 9 | 38 |
| *Nycticebus coucang* | 2 | 10 |
| *Nycticebus pygmaeus* | 4 | 15 |
| *Otolemur crassicaudatus* | 1 | 6 |
| *Otolemur garnettii* | 2 | 10 |
| *Perodicticus potto* | 3 | 13 |
| Total | 113 | 483 |

**Table 2.**
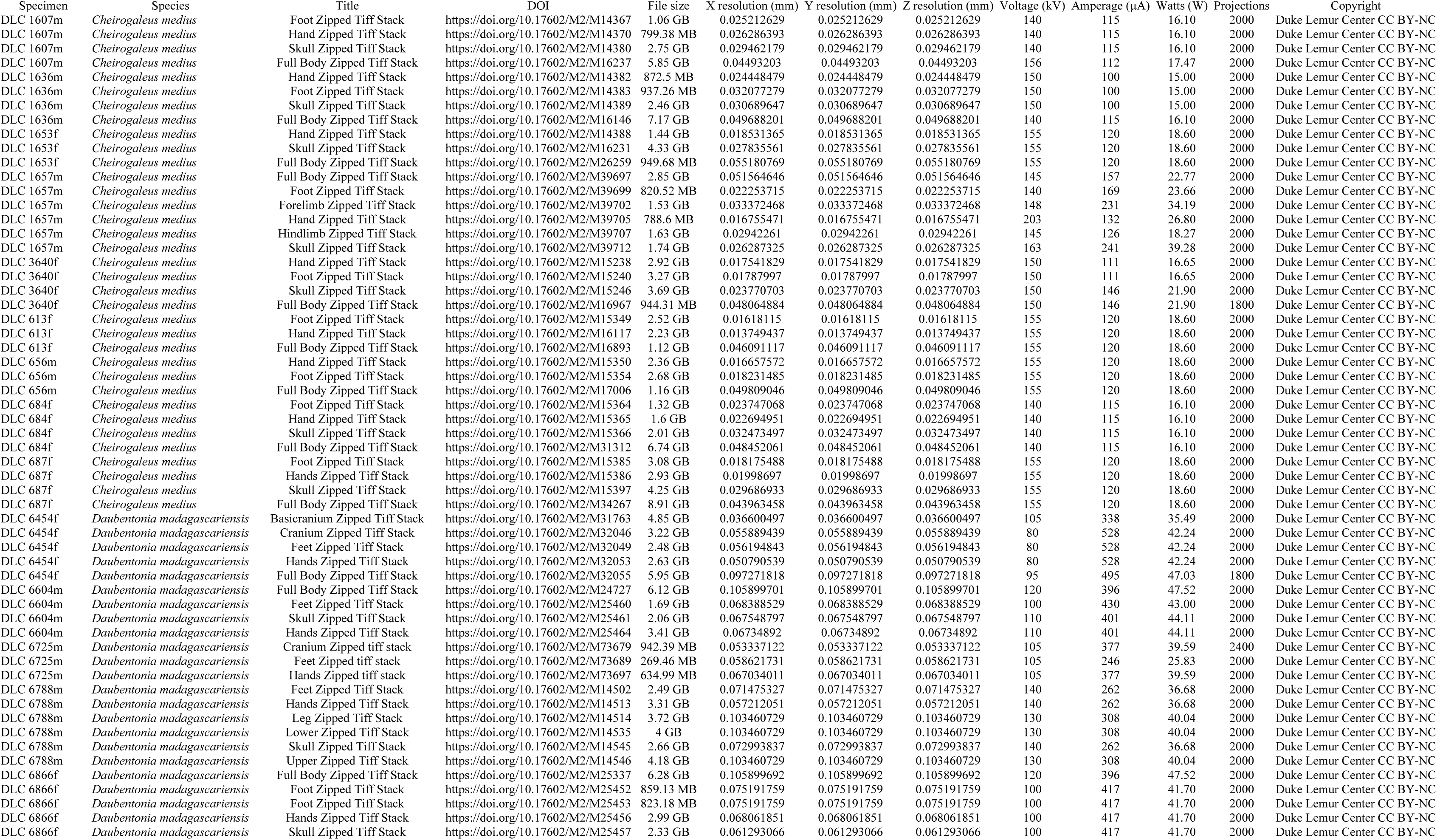
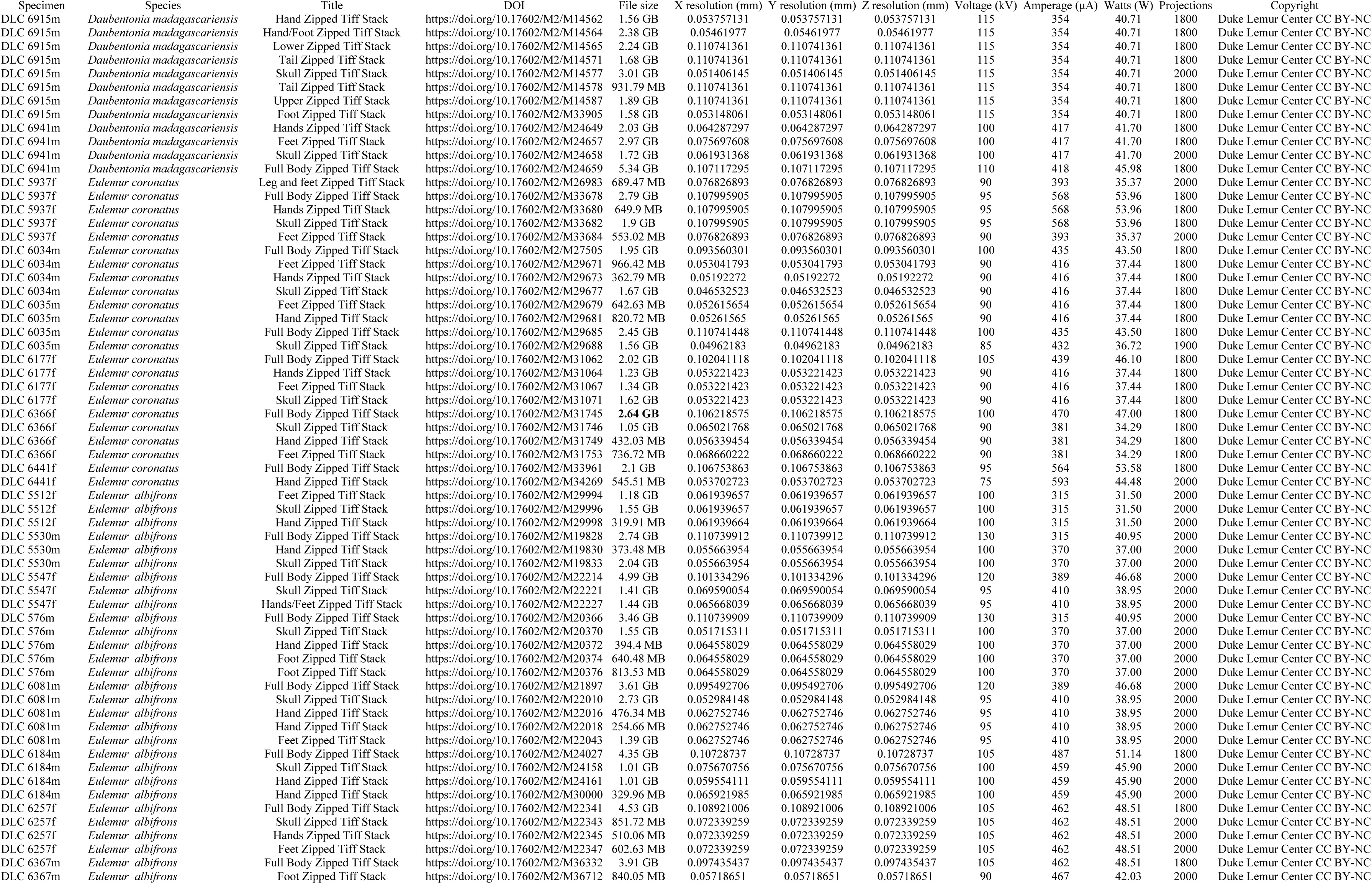
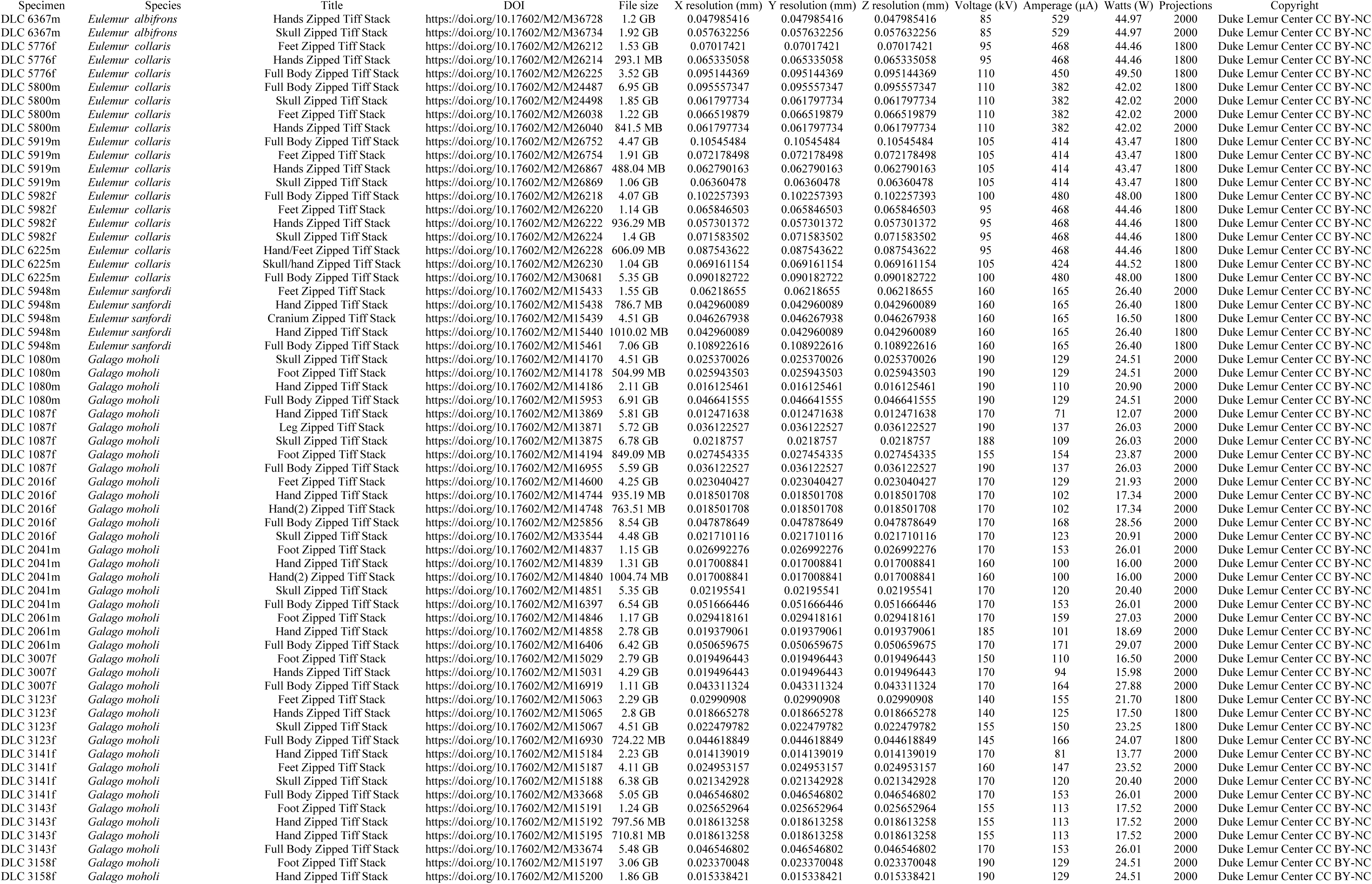
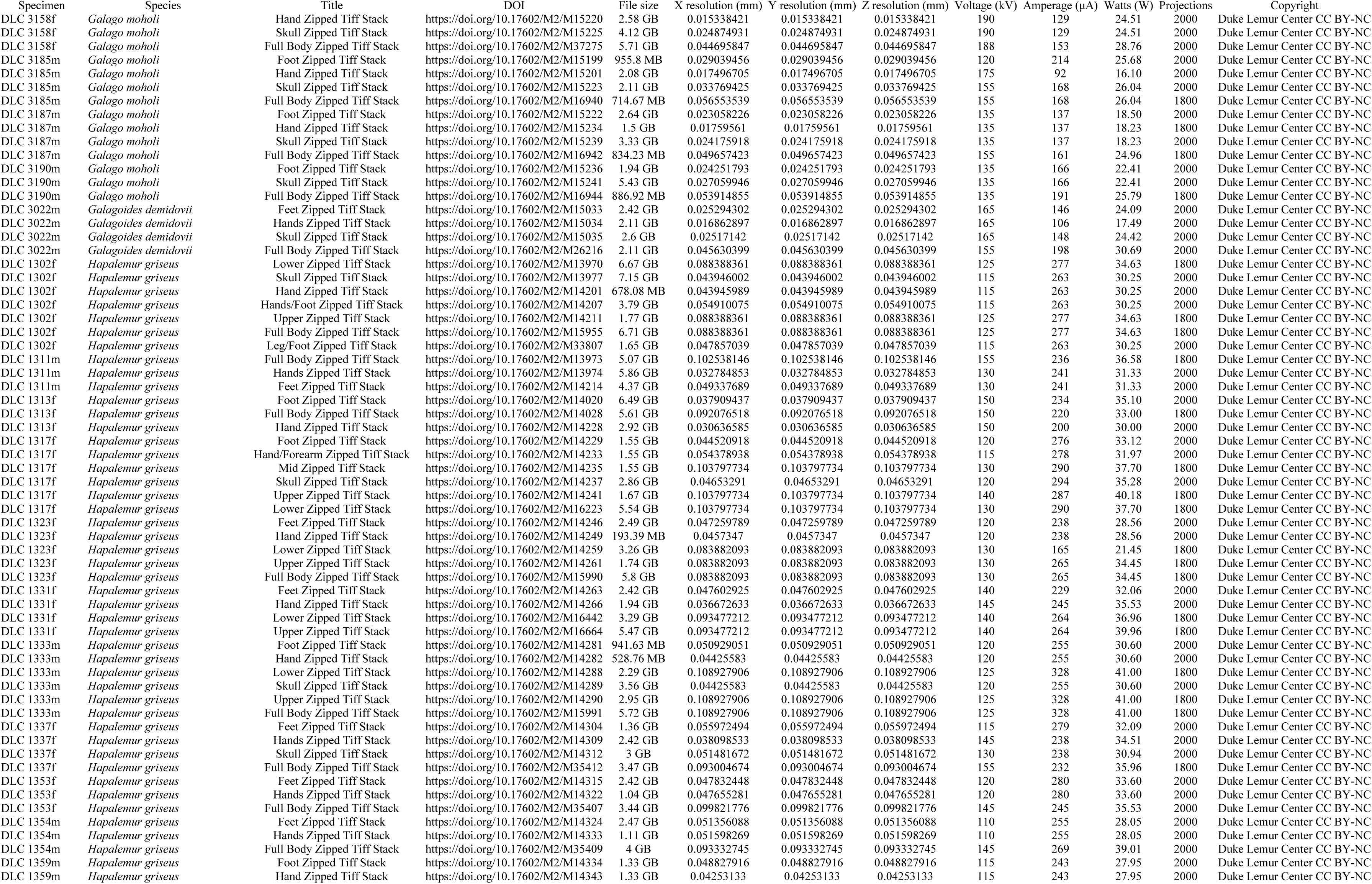
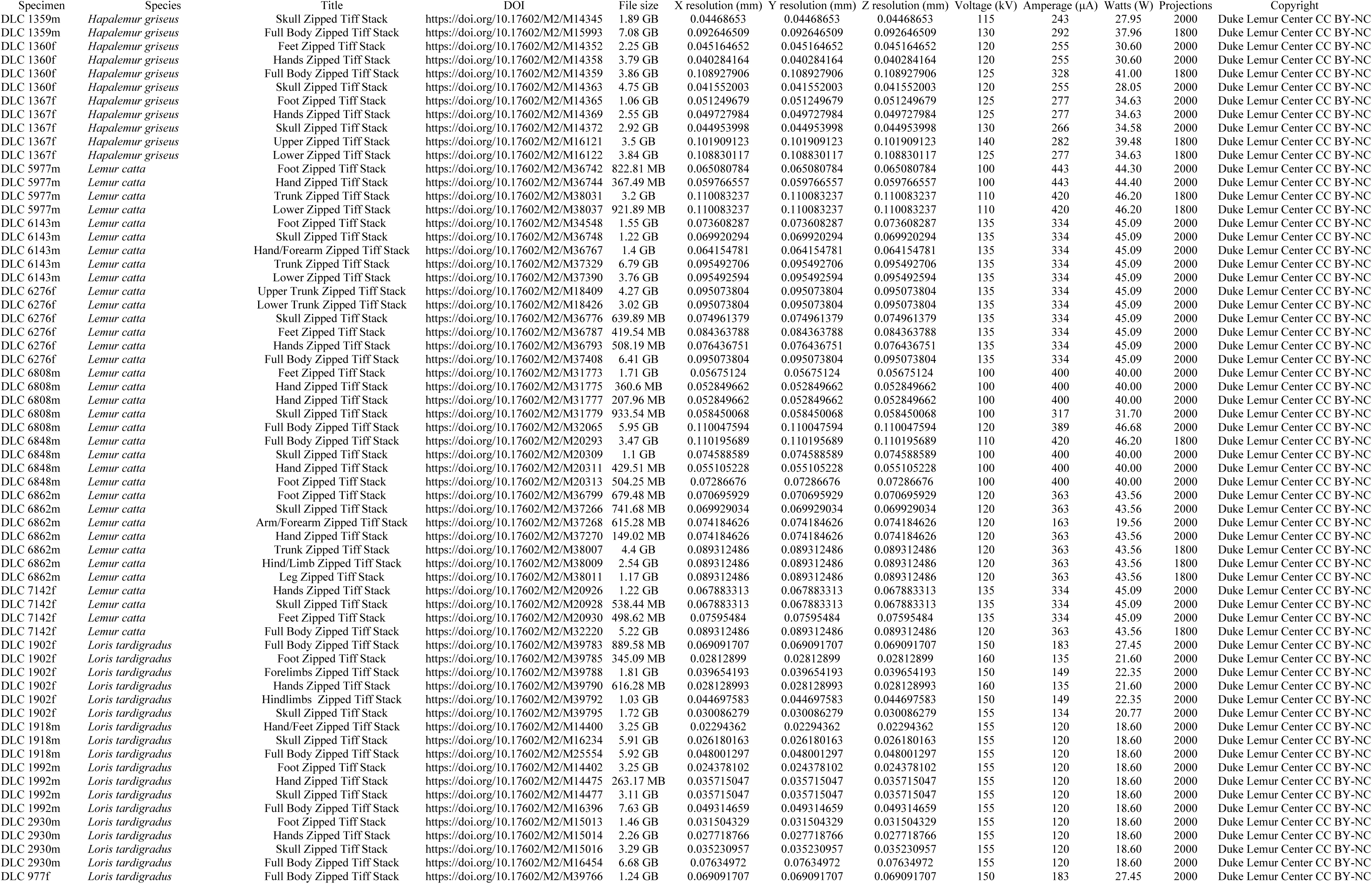
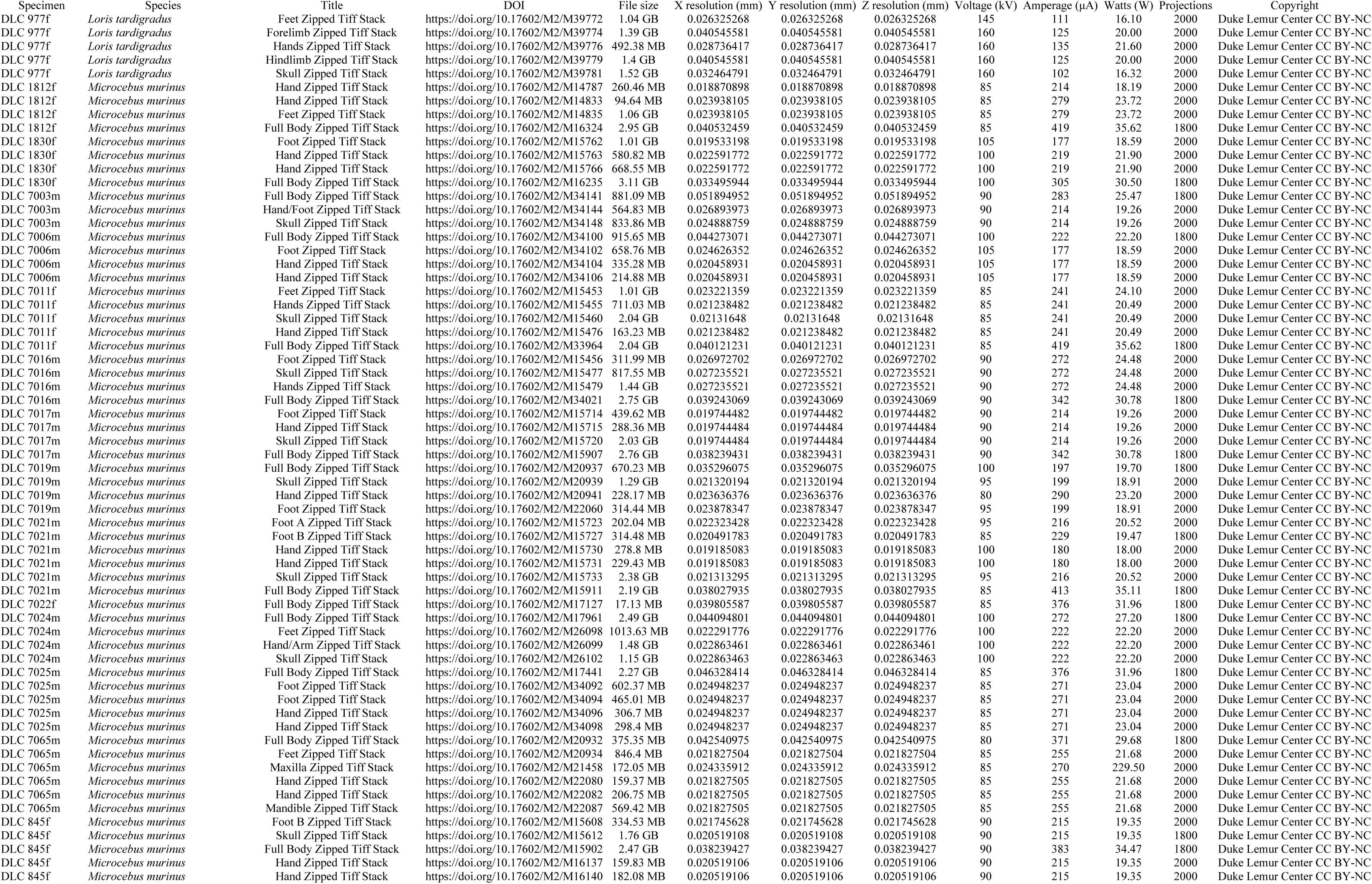
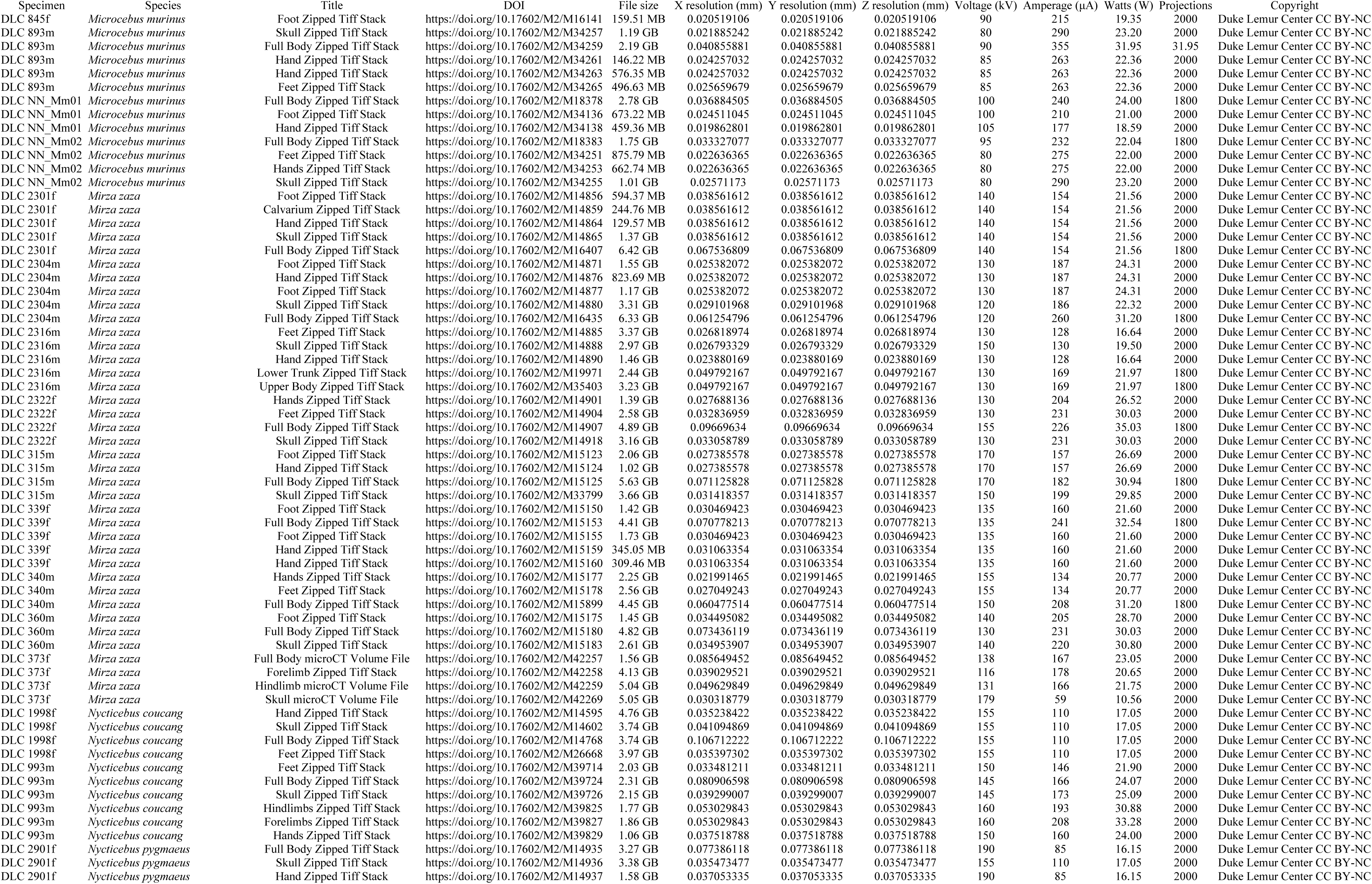
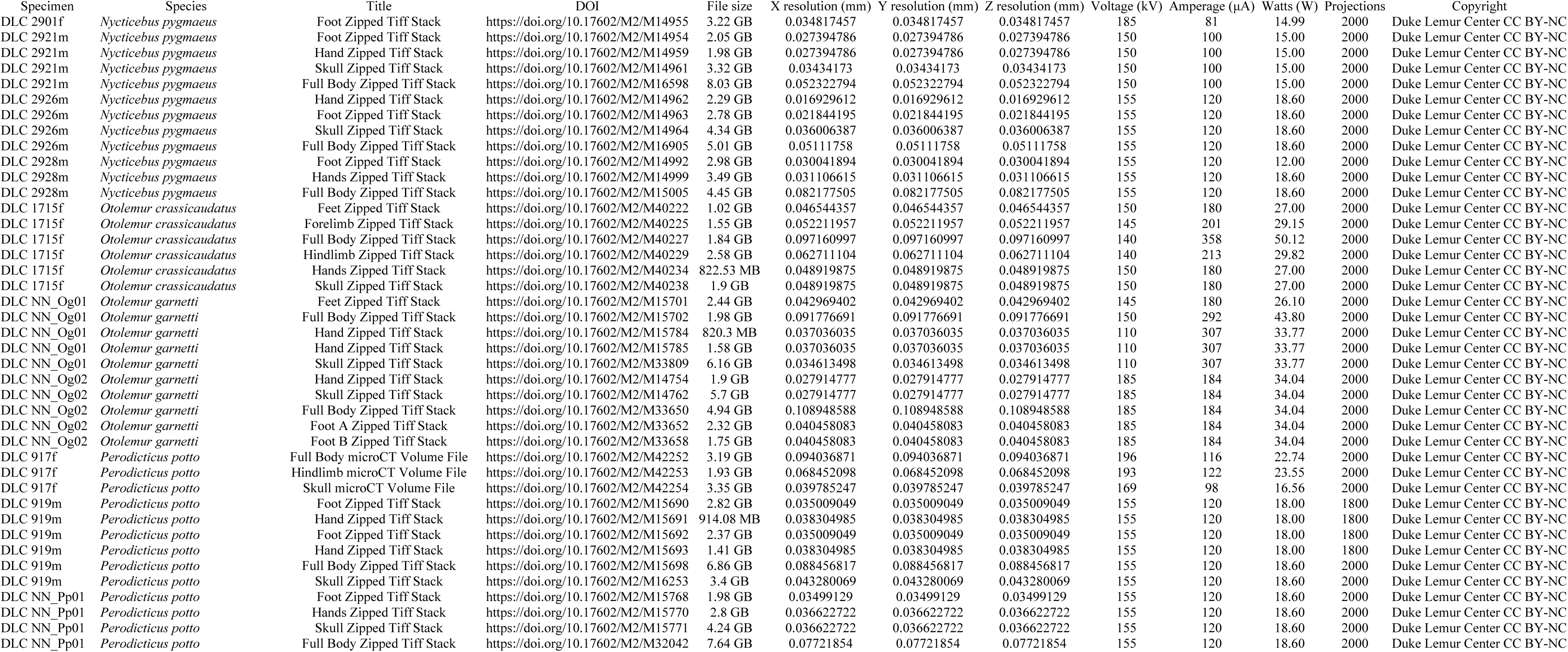
Specimens, scan names, scan parameters, and DOIs for all tiff stacks included in this collection. Abbreviations: mm, millimeters; kV, kilovolts; uA, microamps; GB, gigabytes; MB, megabytes.

### MicroCT scanning

A Nikon XT H 225 ST μCT machine at Duke University’s Shared Materials Instrumentation Facility (SMIF) was used for all scans. This machine has a Perkins Elmer AN1620 X-ray detector panel, which provides a 2000 × 2000 pixel field and a 7.5 frames per second readout. All scans were performed using a Nikon 225kV reflection target with a tungsten anode, which has a focal spot size ranging from 3 to 225 μm depending on the power. No filters were used. Depending on the size and orientation of the specimen, energy settings ranged between 75 – 203 kV and 59 – 593 μA. In order to ensure that each specimen could be used in subsequent research projects, we did not want scanning events to further degrade the specimen through thawing and refreezing. Therefore, we placed non-iodine stained specimens on dry ice in a Styrofoam cooler and conducted the scan with the specimens in the cooler.

The protocol for this project prioritized comprehensiveness and detail. To that end, most specimens were scanned 4 to 8 times and are represented by 4 to 5 volumes: an overview of the full body and separate higher resolution volumes of the skull, hands, and feet (Figure 2). Full body overviews are usually composites created by stitching together tiff stacks from multiple scanning events (detailed further below).

**Fig 2.**
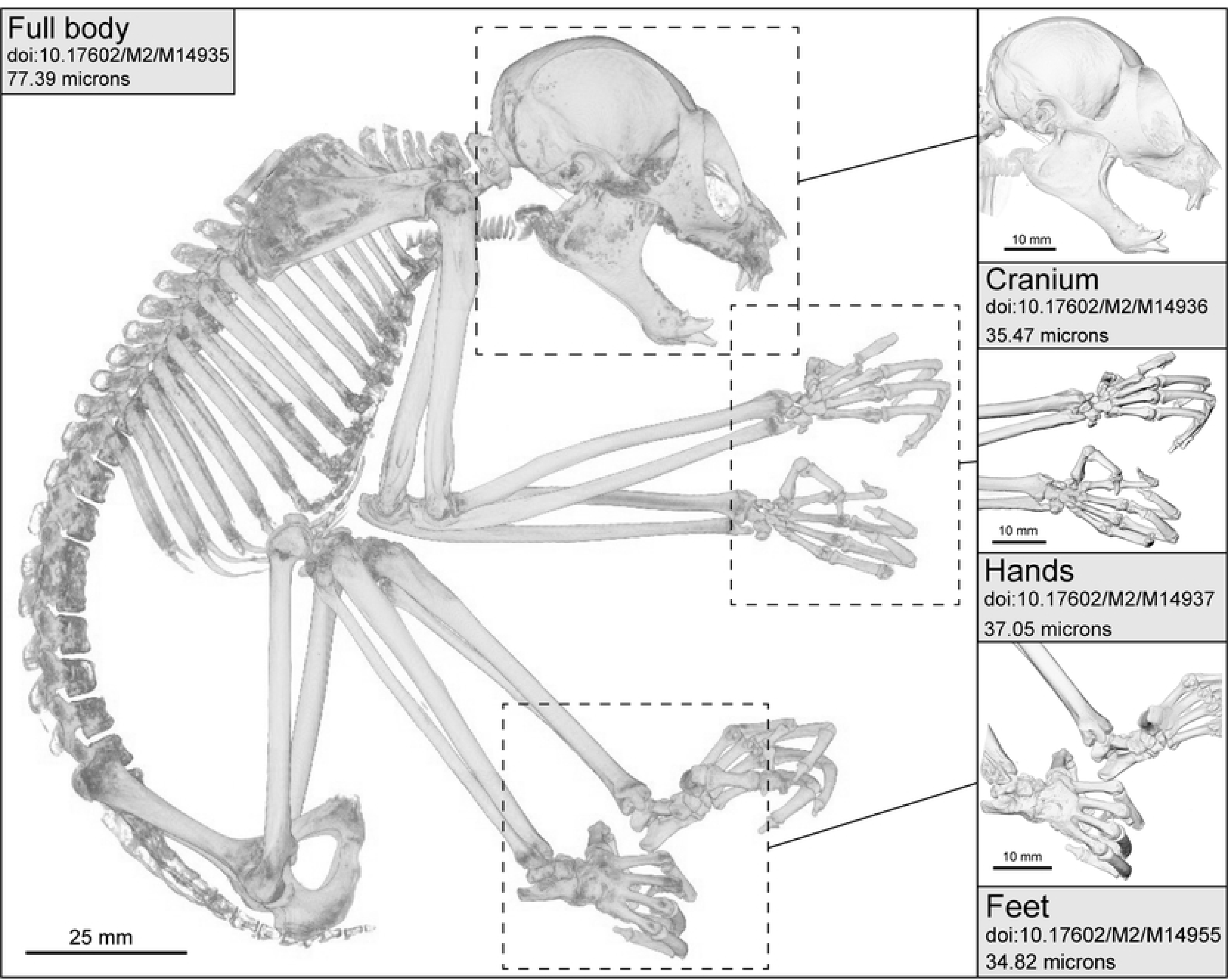
Volume rendering of Nycticebus coucang (DLC 2901f) showing scanning protocol for strepsirrhine cadavers. Boxes outline regions of increased anatomical complexity that were scanned at higher resolutions separately. To reduce noise, the threshold for grey values is lower than optimal threshold, rendering less dense bone transparent.

### Iodine staining

The soft tissue anatomy of two lemur and five loris specimens was visualized in the CT-scans using iodine as a contrasting agent [12]. Figure 3 provides an example image from an individual scanned after the staining process. These specimens were first thawed and then fixed in formalin (Carolina Biological), then stained in a 7% solution of Lugol’s Iodine (Carolina Biological) for six weeks prior to scanning.

**Fig 3.**
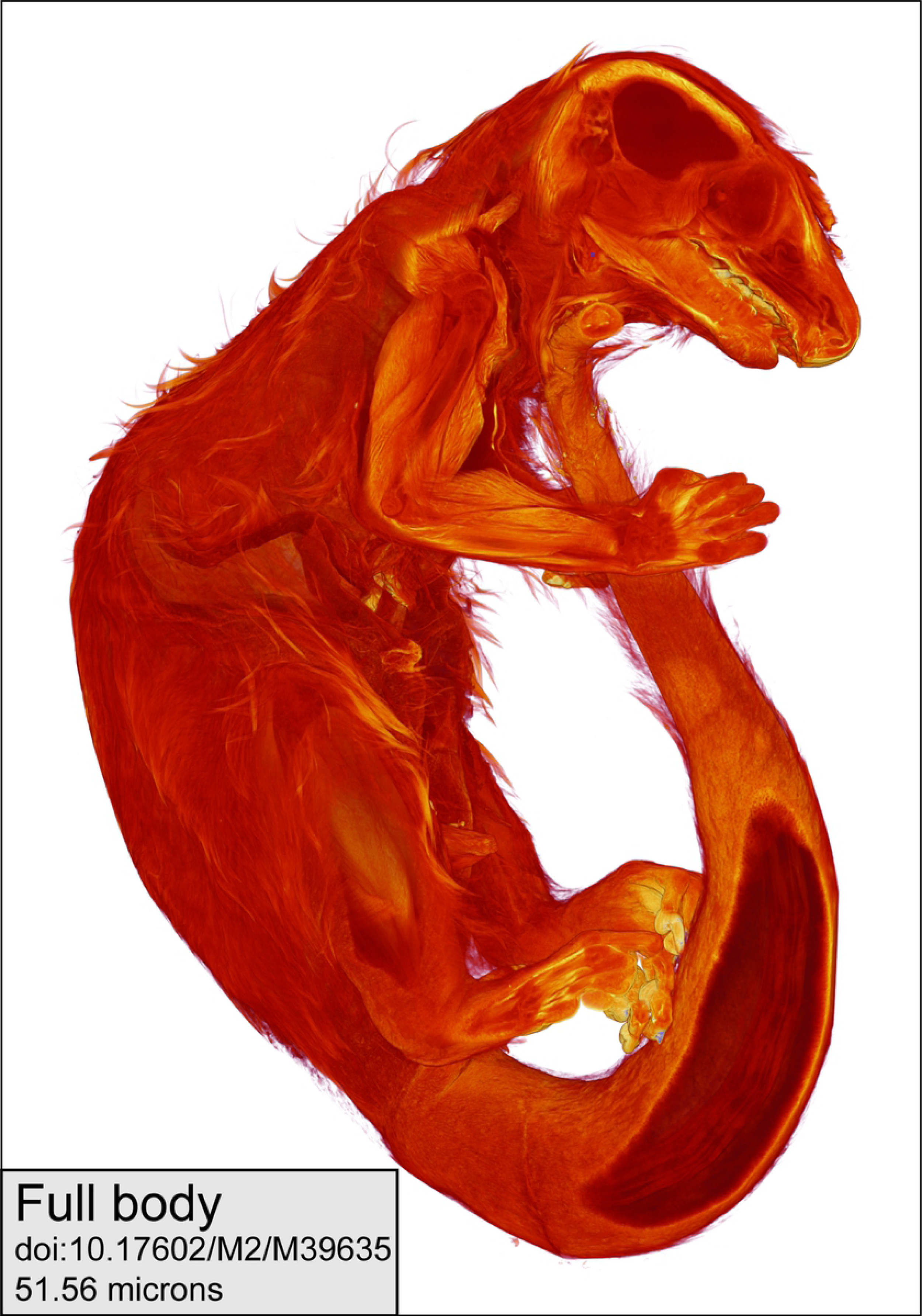
Volume rendering of iodine-stained Cheirogaleus medius (DLC 1657m) using Avizo software. A clipping plane in the software digitally slices through the fur and skin to show stained tissues underneath.

### Post-processing and stitching

X-ray projections were reconstructed as 3D volumes using Nikon software. Volumes were saved as 16-bit Tagged Image File Format (.tiff) stacks. High resolutions scans were compressed with 7-Zip and uploaded to MorphoSource.

If the size of the specimen prevented the full body overview from being done in a single scan, the overview was created by conducting a series of overlapping scans with a shared centre of rotation. This process was easy to accomplish when the specimen was in an extended posture and each scan was oriented along the vertical y-axis (Figure 4a), provided the specimen was not larger than the vertical travel distance permitted by the scanner’s dimensions. However, larger specimens were often flexed into C-shapes that exceeded the dimensions of the detector, even at the lowest magnification. In these cases, scans were conducted as a series of overlapping “panels” with two separate vertical axes (Figure 4b).

**Fig 4.**
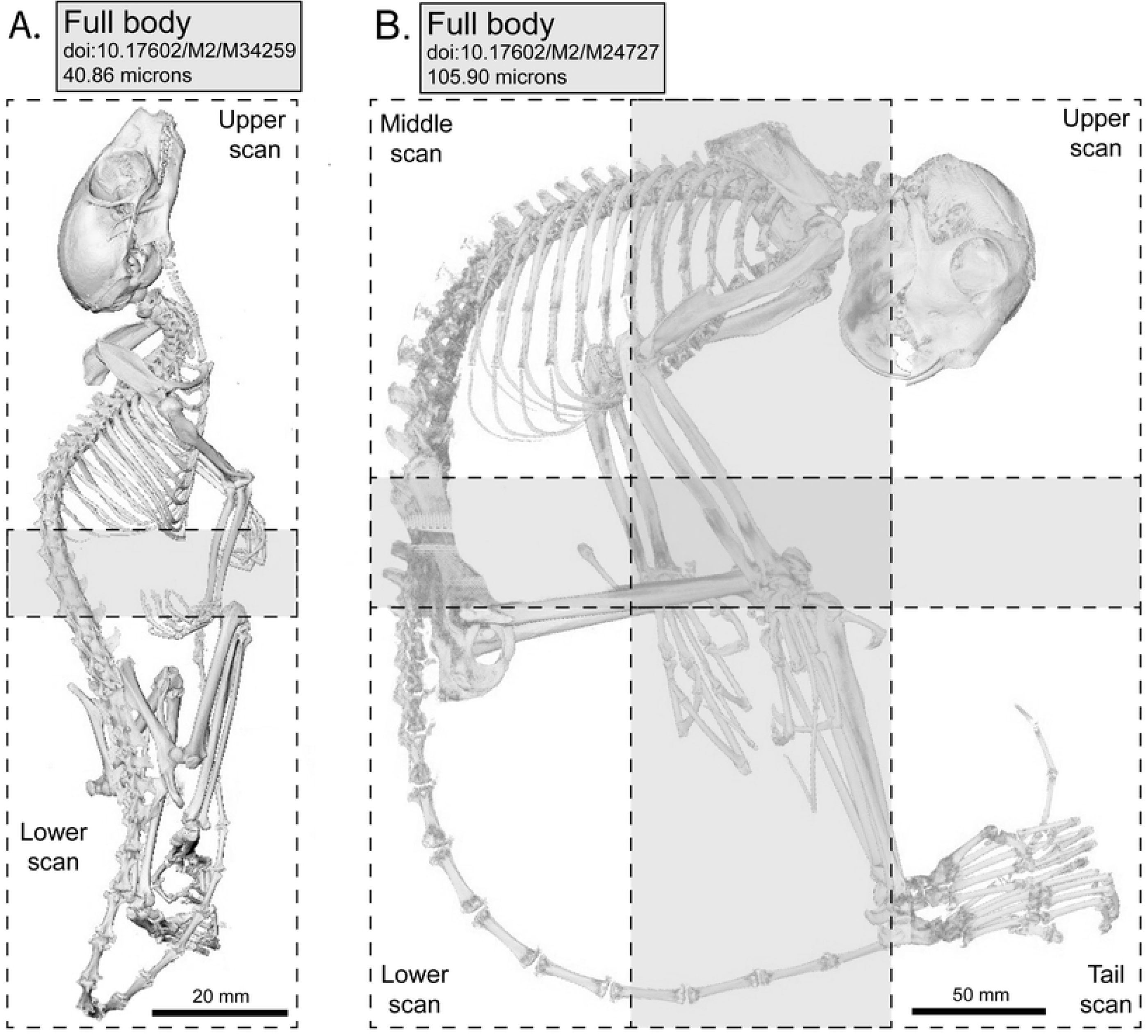
Examples of stitched composite scans. A) composite scan of *Microcebus murinus* (DLC 893m) stitched along a single vertical axis; B) composite scan of *Daubentonia madagascariensis* (DLC 6604m) stitched along two vertical axes. Boxes indicate separate microCT scans, grey areas boxes indicate areas of scan overlap.

To generate full body composites, overlapping scans were stitched together in ImageJ [13] using the 3D Stitching plug-in [14]. As composites of two or more scanning events, the full body overviews are very large volumes. When composite overviews were extremely large (rendering subsequent processing too computationally demanding), we chose to upload the overlapping scans separately. If there are elements partially represented across two scans, these scans can be stitched together by researchers after download.

### MicroCT error study

As discussed by Copes et al. [9], μCT scanners in academic instrumentation facilities accommodate a wide range of users with varying demands for scanning parameters (i.e., detector and stage settings, target type, beam settings). The flexibility required of μCT scanners in academic contexts stands in contrast to industrial or metrology-specific machines, which can be calibrated to maintain a degree of minimum error within a particular set of scan parameters. For scanners in academic instrumentation facilities, accuracy is determined by the initial installation settings and subsequent maintenance, with the assumption that measurement error is around 1%.

Here we expand the calibration study of Copes et al. [9] for SMIF’s Nikon XT H 225 ST μCT machine. To determine the accuracy of the scanner, three different standard spheres of known diameters (3.175mm, 6.35mm, and 12.7mm; all +/- 1.0 um tolerance) were scanned at a range of voxel resolutions. The 3.175mm and 6.35mm spheres was scanned at 5, 6, 7, 8, 9, 10, 15, and 20 µm per voxel with the detector fixed at its farthest position in the chamber. The 12.7mm sphere was scanned at 10, 20, 30, 40, 50, 60, 70, 80, 90, and 100 µm per voxel with the detector in the same position. Each scan was collected at 175kV, 86µA (15W), 354ms, 2000 projections, 1 frame per projection, and without a filter. Nikon’s proprietary CT Pro 3D and CT Agent software reconstructed the projection data into a volume data file, which was then opened in VG Studio Max 2.2.

In VG Studio, an automatic surface determination was applied to the spheres, >20 fit points were placed on the surface, and an idealized sphere was fit to these points. Diameters produced from this measurement were recorded and compared to the reported diameter of the spheres. The relative percentage error (RE%) was calculated as the difference between the measured (MD) and reported diameters (RD), divided by the measured diameter (RE% = (MD-RD)/MD*100). Given that we are evaluating the same μCT machine, we expect relative % error to be similar to the <0.2% reported by Copes et al. [9].

## Results

### MicroCT error study

Table 3 reports error values for all three standard spheres, and error is plotted against voxel resolution in Figure 5. For each calibration sphere, we found less than 0.3% error at all resolution levels, with a majority of scans demonstrating error levels below 0.1%.

**Table 3.**
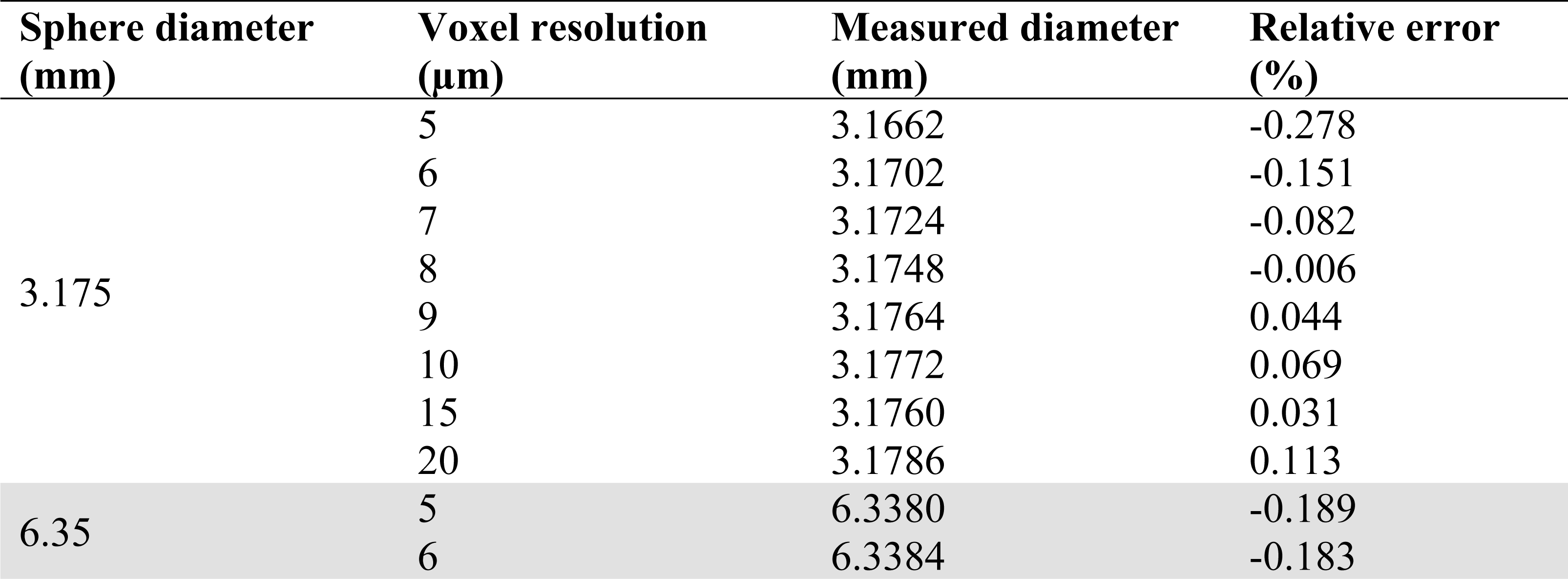
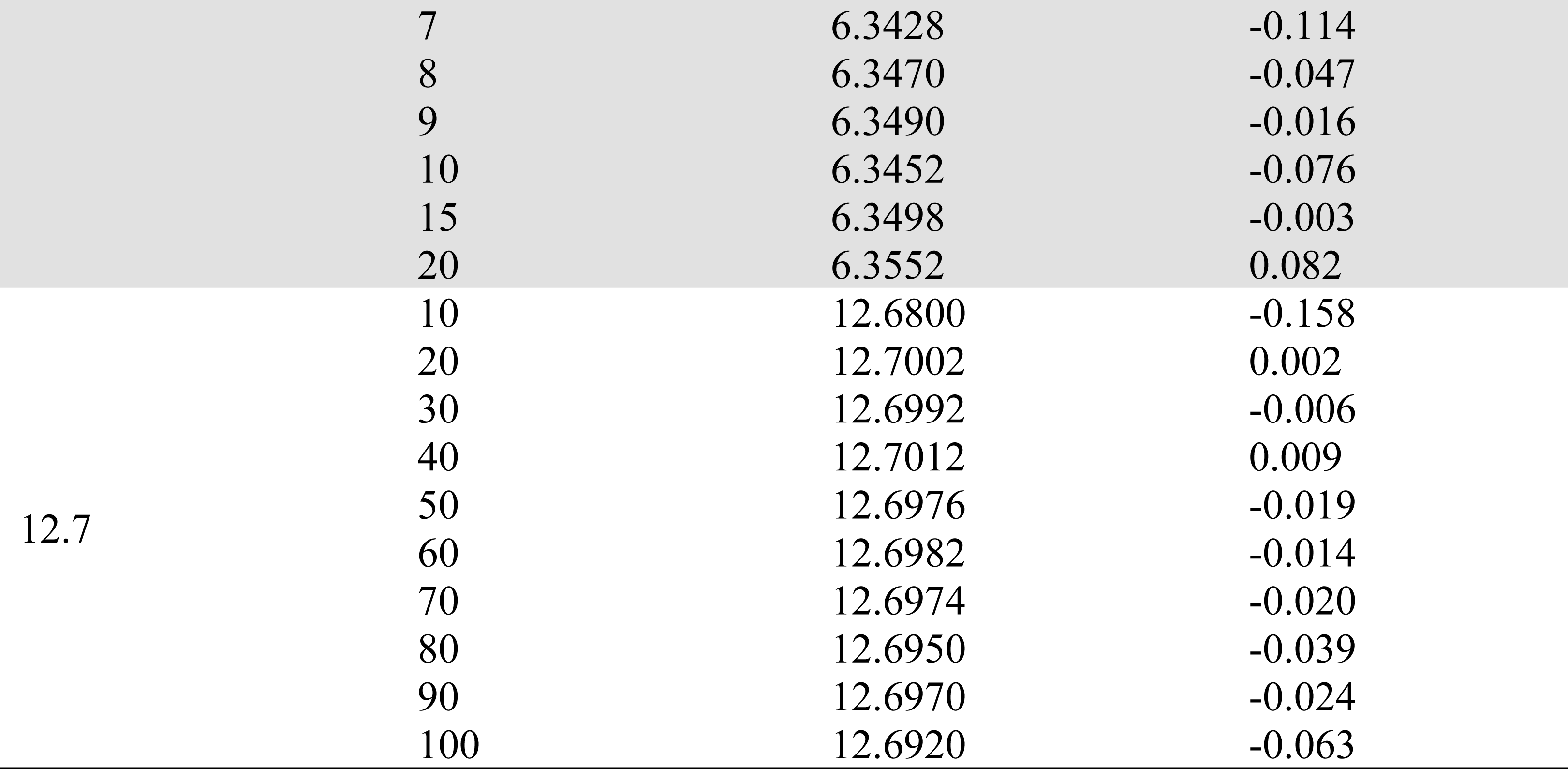
Relative error at different voxel resolutions for three calibration spheres in the Nikon X-Tek XHT 225 ST scanner at Duke University’s Shared Materials and Instrumentation Facility.

**Fig 5.**
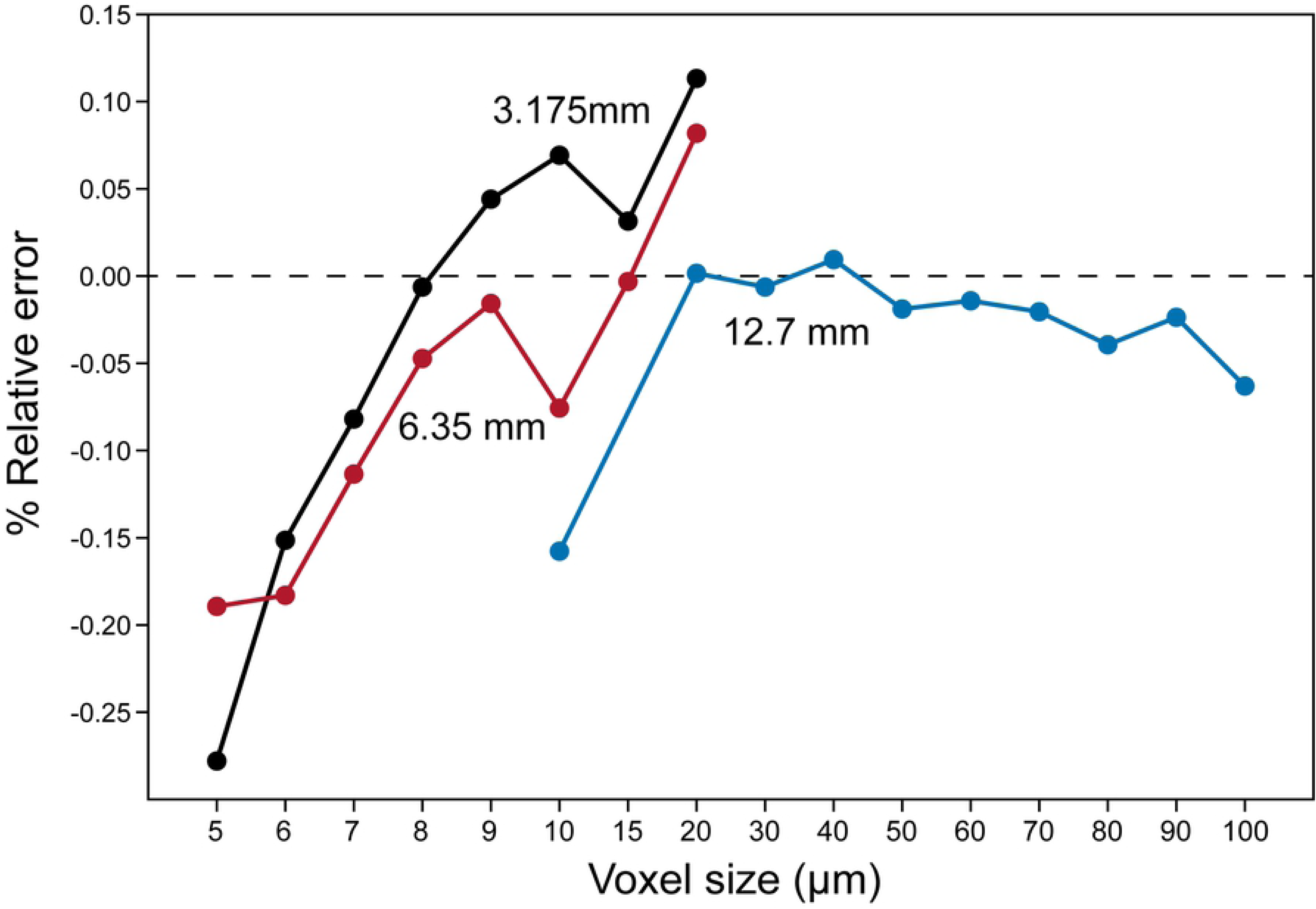
Relative error at different voxel resolutions for three calibration spheres (diameters of 3.175 mm, 6.35 mm, and 12.7 mm) in the Nikon XT H 225 ST μCT scanner at Duke University’s Shared Materials Instrumentation Facility.

### Data records and availability

The μCT scans and associated metadata (Table 2) presented in this project are available through MorphoSource (http://www.morphosource.org), a free-to-use online repository designed to accommodate 3D data and its derivatives, including 3D surface renderings and other digital imagery. MorphoSource was created to address increasing demand for deposition and archiving of 3D digital data, and provides the necessary infrastructure to host, share, and manage 3D data for several different user types—from individual researchers to museum curators—allowing data authors and institutions to benefit from subsequent data use by third parties. Network storage provided by Duke University’s data infrastructure with servers housed in multiple locations.

Data files of the current project are available open access. Data files are copyrighted by the DLC and can be downloaded and re-used for non-commercial purposes (Creative Commons copyright license CC-BY NC). As of June 12th, this project has been viewed more than 43,000 times and more than 1200 data files have been downloaded. Because scanning efforts of the DLC cadaver collection are ongoing, this project will continue to grow to include new specimens and new species.

## Discussion

### Structure of the digital collection

We chose MorphoSource to host this collection because it offers several utilities for data authors and institutions. First, MorphoSource is structured to function as both an online platform for collaborative research and as an institutional repository for 3D digital data, with high ease-of-use when transitioning between these two roles. For this project, we began to upload scans in 2016 and, for the next two years, data were shared privately with collaborators at multiple institutions. When the project matured, data were made searchable and downloadable to all MorphoSource users, signalling the project’s transition to a publicly facing data collection. In providing this dual functionality, MorphoSource has utility for data authors during all stages of 3D data collection.

Second, MorphoSource provides data authors and institutions multiple ways to track and summarize data usage, including summary reports of data usage as well as the ability to establish citable digital object identifiers (DOIs) for individual scans and derivative data. These tools provide metrics of the value of the collection and help provide professional benefits for researchers undertaking time-intensive data collection efforts. Table 2 was generated from the summary report of this project.

Third, while data collections can be made publicly available through MorphoSource, the platform allows data authors or institutions the option of managing user access through a data request system. This functionality, along with the ability to clearly delineate copyright restrictions and other terms of use, reduces concerns that data will be used in ways that are inconsistent with institutional goals.

Data files on MorphoSource are organized within a three-level hierarchy: specimens > media groups > media files. A fourth organizational unit, the MorphoSource project, exists at the same hierarchical level as the specimen (i.e., specimens are not contained within projects, although media records of a specimen might be). Projects are a collection of media records determined by the user and serve several purposes, including sharing multiple data files with collaborators easily, developing special collections of high public interest (e.g., K-12 Anthropology Teaching Collection, MorphoSource Project P158), or curating institutional holdings (such as this project).

On MorphoSource, specimens are digital representations of physical specimens stored in museums or other repositories. As researchers upload new data, they increase the comprehensive sampling of MorphoSource specimens. When new specimen records are created, they can be linked with the institution’s metadata through iDigBio (http://www.idigbio.org).

Media groups are nested within specimens and generally represent discrete scanning or other digitization events. Metadata of the digitization event (including scan parameters, calibrations, and funding sources) are attached to media groups.

Media files are nested within media groups and represent raw data (such as tiff stacks) and derivative data (such as surface renderings). When media files are made publicly available or “published”, they can be assigned DOIs, which function as permanent and direct links to the data as well as references for data citation in subsequent studies. Currently, the DLC cadaver project contains media files tagged by 1329 digital object identifiers, including the 483 raw tiff stacks presented in this study and 374 derivative surface files. Further detail on the structural organization and associated metadata of MorphoSource can be found in Boyer et al. [5].

### Computer requirements

By current computer standards, these μCT scans are large files. Tiff stacks can range from 1 to 13 GBs and the stitched full body composites are particularly large. System performance depends primarily on four hardware components: the graphics processing unit (GPU), the central processing unit (CPU), the amount of random-access memory (RAM), and the hard drive. For direct volume renderings or 3D surface visualization, the system should have a high-end graphics card (minimally a DDR5 memory interface with more than 2 GB RAM). Sufficient RAM is critical for 3D visualization and analyses; we follow Copes et al. (2016) in recommending computers have RAM that is at least twice as large as the largest tiff stack. Image processing speed relies on the CPU, so processors with high clock speeds (greater than 3 gigahertz) are recommended. Finally, for uploading datasets quickly, solid state drives (SSD) provide much faster reading and writing speeds than traditional hard disk drives (HDD).

Other recommendations to improve performance are a 64-bit operating system and multiple processing cores. Specific 3D visualization programs (listed below) may have their own set of system recommendations, so researchers are encouraged to evaluate the match between their preferred software and available workstation.

### Data manipulation

When downloaded from MorphoSource, these μCT scans will first need to be decompressed using commercially available programs such as WinZip or 7-Zip. After decompression, we recommend that users then open tiff stacks with 2D image processing software such as Fiji [15] or ImageJ [13], as these programs permit some volume editing but require less memory than industrial volume visualization and analysis software. In Fiji or ImageJ, the user can easily adjust grey values to enhance contrast and crop the original μCT volume to regions of interest. Edited tiff stacks can be saved as new image sequences that require less memory to open and manipulate. For 3D visualization and analyses, there are several commercially available programs (e.g., Avizo, Amira, Dragonfly, Mimix, Osirix, Spiers, and VG Studio Max) as well as freeware (e.g., Slicer3D).

### Citation of scans

Usage terms are outlined in the MorphoSource user agreement that accompanies each download from the website. Subsequent publications that make use of these scans should include 1) a citation of this study; 2) a list or table of DOIs for each scan used; and 3) a statement of access accompanying the DOI list or in the acknowledgments. This statement should read “The Duke Lemur Center provided access to these data, originally appearing in Yapuncich et al. (2019), the collection and archiving of which was funded by NSF BCS 1540421 and NSF BCS 1552848. The files were downloaded from www.MorphoSource.org, Duke University.” A similar statement is included in the accompanying scan metadata.

Although not required by the usage agreement, we would urge researchers who conduct additional processing of these scans (e.g., generate surface files of elements of interest) to upload derivative files to MorphoSource. While we recognize that uploading derivatives can be a time-intensive process, we feel it is a critical component of data sharing and subsequent research. To facilitate this process, bulk upload options have been developed and are available through MorphoSource. Finally, we encourage researchers to contact the DLC research manager to obtain a DLC publication number when manuscripts using these data have been accepted for publication.

## Acknowledgments

The authors thank Kay Welser, David Brewer, Anne Yoder, and Greg Dye at the Duke Lemur Center for facilitating specimen loans, Mackenzie Shepherd for helping upload scans to MorphoSource, and Liza Shapiro for access to the specimens for iodine-staining. This work was performed in part at the Duke University Shared Materials Instrumentation Facility (SMIF), a member of the North Carolina Research Triangle Nanotechnology Network (RTNN), which is supported by the National Science Foundation (Grant ECCS-1542015) as part of the National Nanotechnology Coordinated Infrastructure (NNCI).

## Author contributions

G.S.Y. scanned specimens, uploaded them to MorphoSource, and helped write the paper. A.D.K. stained and scanned specimens and helped write the paper. D.M.G. processed scans and uploaded them to MorphoSource. J.T.G. conducted calibration study of the microCT scanner and helped write the paper. E.E. facilitated specimen access and helped write the paper. D.M.B. created and manages MorphoSource, facilitated the uploading process, and helped write the paper.

## References

1. Schwitzer C, Mittermeier RA, Johnson SE, Donati G, Irwin M, Peacock H, Ratsimbazafy J, Razafindramanana J, Louis EE, Chikhi L, Colquhoun IC. Averting lemur extinctions amid Madagascar’s political crisis. Science. 2014; 343:842–843.

2. Schwitzer C, Mittermeier RA, Davies N, Johnson S, Ratsimbazafy J, Razafindramanana J, Louis Jr EE, Rajaobelina S. Lemurs of Madagascar: A strategy for their conservation 2013–2016. Bristol, UK: IUCN SSC Primate Specialist Group, Bristol Conservation and Science Foundation, and Conservation International; 2013.

3. Yapuncich GS. Body mass prediction from dental and postcranial measurements in primates and their nearest relatives. Ph. D. Dissertation, Duke University. 2017.

4. Kemp AD. Primate binocular vision: adaptive significance in grasping and locomotion. Ph. D. Dissertation, University of Texas, Austin. 2019.

5. Boyer DM, Gunnell GF, Kaufman S, McGeary TM. Morphosource: Archiving and sharing 3-d digital specimen data. Paleontol Soc Pap. 2016; 22:157–181.

6. Mittermeier RA, Fleagle JG. A primate distribution program to end wastage of sacrificed specimens. Lab Prim News. 1973; 12: 1–3.

7. Gordon AD, Marcus E, Wood B. Great ape skeletal collections: making the most of scarce and irreplaceable resources in the digital age. Am J Phys Anthropol. 2013; 152:2–32.

8. Adams JW, Olah A, McCurry MR, Potze S. Surface model and tomographic archive of fossil primate and other mammal holotype and paratype specimens of the Ditsong National Museum of Natural History, Pretoria, South Africa. PLoS One. 2015; 10:e0139800.

9. Copes LE, Lucas LM, Thostenson JO, Hoekstra HE, Boyer DM. A collection of non-human primate computed tomography scans housed in MorphoSource, a repository for 3D data. Sci Data. 2016; 3:160001.

10. Shi JJ, Westeen EP, Rabosky DL. Digitizing extant bat diversity: An open-access repository of 3D μCT-scanned skulls for research and education. PLoS One. 2018; 13:e0203022.

11. Zehr SM, Roach RG, Haring D, Taylor J, Cameron FH, Yoder AD. Life history profiles for 27 strepsirrhine primate taxa generated using captive data from the Duke Lemur Center. Sci Data. 2014; 1:140019.

12. Gignac PM, Kley NJ, Clarke JA, Colbert MW, Morhardt AC, Cerio D, Cost IN, Cox PG, Daza JD, Early CM, Echols MS. Diffusible iodine-based contrast-enhanced computed tomography (diceCT): an emerging tool for rapid, high-resolution, 3-D imaging of metazoan soft tissues. J Anat. 2016; 228:889–909.

13. Schneider CA, Rasband WS, Eliceiri KW. NIH Image to ImageJ: 25 years of image analysis. Nature Methods. 2012; 9:671.

14. Preibisch S, Saalfeld S, Tomancak P. Globally optimal stitching of tiled 3D microscopic image acquisitions. Bioinformatics. 2009; 25:1463–1465.

15. Schindelin J, Arganda-Carreras I, Frise E, Kaynig V, Longair M, Pietzsch T, Preibisch S, Rueden C, Saalfeld S, Schmid B, Tinevez JY. Fiji: an open-source platform for biological-image analysis. Nature Methods. 2012; 9:676.

16. Arnold C, Matthews LJ, Nunn CL. The 10kTrees website: a new online resource for primate phylogeny. Evol Anthropol. 2010; 19:114–118.

